# Distribution of genetic diversity reveals colonization and philopatry of the loggerhead sea turtles across geographic scales

**DOI:** 10.1101/2020.01.23.916866

**Authors:** Miguel Baltazar-Soares, Juliana L. Klein, Sandra M. Correia, Thomas Reischig, Amoros Albert Taxonera, Roque Silvana Monteiro, Leno Dos Passos, Durão Jandira, Pina Lomba João, Herculano Dinis, Sahmorie J.K. Cameron, Victor A. Stiebens, Christophe Eizaguirre

## Abstract

**Aim:** Understanding the processes that underlie the distribution of genetic diversity in endangered species is a goal of modern conservation biology. Specifically, how population structure affects genetic diversity and contributes to a species’ adaptive potential remain elusive. The loggerhead sea turtle (*Caretta caretta*) faces multiple conservation challenges due to its migratory nature and philopatric behaviour.

**Locations:** Atlantic Ocean, Cabo Verde, island of Boavista

**Methods:** Here, using 4207 mtDNA sequences, we analysed the colonisation patterns and distribution of genetic diversity within a major ocean basin (the Atlantic), a regional rookery (Cabo Verde Archipelago) and a local island (Island of Boavista, Cabo Verde).

**Results:** Hypothesis-driven population genetic models suggest the colonization of the Atlantic has occurred in two distinct waves, each corresponding to major mtDNA lineages. We propose the oldest lineage entered the basin via the isthmus of Panama and sequentially established aggregations in Brazil, Cabo Verde and in the area of USA and Mexico. The second lineage entered the Atlantic via the Cape of Good Hope, establishing colonies in the Mediterranean Sea, and from then on, re-colonized the already existing rookeries of the Atlantic. At the Cabo Verde level, we reveal an asymmetric gene flow maintaining links across nesting groups despite significant genetic structure amongst nesting groups. This structure stems from female philopatric behaviour which could further be detected by weak but significant structure amongst beaches separated by only a few kilometres on the island of Boavista.

**Main conclusion:** To explore demographic processes at diverse geographic scales improves understanding the complex evolutionary history of highly migratory philopatric species. Unveiling the past facilitates the design of conservation programmes targeting the right management scale to maintain a species’ adaptive potential and putative response to human-induced selection.

## Introduction

The distribution of species is shaped by environmental variation acting both at macro and micro evolutionary scales (Ezard et al., 2011). Nowadays the distribution of biodiversity reflects specie’s evolutionary history as well as eco-evolutionary dynamics within and across systems (Brunner et al., 2019). Because environmental factors (e.g. (Perry et al., 2005)) and past events of colonization (e.g. (Carotenuto et al., 2016)) leave genetic signatures, characterizing the distribution of genetic diversity sheds light on the mechanisms underlying the species and populations’ distributions e.g. (Loveless & Hamrick, 1984; Ellegren & Galtier, 2016). Understanding those mechanisms is crucial for the implementation of management and conservation measures to maintain species’ adaptive potential (Frankham et al., 2002; Eizaguirre & Baltazar-Soares, 2014). The recent years have seen an increasing number of tools available to performed model-based inferences in evolutionary genetics (Csillery et al., 2010). Inferences are made over likelihood estimates of a certain set of pre-defined parameters constituting different evolutionary scenarios (Beaumont et al., 2010). Likelihood estimates can be obtained, for example, with exact calculations and usually use coalescent theory and Bayesian statistics to simulate phylogenies with Markov Chain Monte Carlo (MCMC) samplers (Drummond et al., 2005). Alternatively, likelihood estimates can be approximated with Approximate Bayesian Computation (ABC) mehtods which emmerged as an alternative that derives likelihood estimates through comparisons of summary statistics of simulated datasets and those obtained with observed data (Csillery et al., 2010). Being less demanding computationally, ABC facilitates hypothesis-testing to explain phylogeographic patterns that would be otherwise challenging to explore through computing exact likelihood estimates. For marine species, model-based inferences have contributed, for example, to understand oceanic divergence of humpback whales or the post glacial distribution of blacknose sharks (Jackson et al., 2014; Portnoy et al., 2014), revealing a promising tool to investigate the complex demographic patterns of philopatric and highly migratory species. Philopatry, the tendency of an organism to return to its home area or natal site to reproduce (Mayr, 1963; Greenwood, 1980), impacts the genetic structure of species, forming groups of individuals of similar matrilineage (Mourier & Planes, 2013; Stiebens et al., 2013). This strategy is common in the aquatic realm [e.g. salmonids (Dittman & Quinn, 1996), cetaceans (Engelhaupt et al., 2009), sharks (Hueter et al., 2005) and turtles (Bowen et al., 2005; Carreras et al., 2007; Lee, 2008)]. Particularly, sea turtles are capable of homing on a scale of a few kilometres (Lee et al., 2018). Upon hatching, neonate enter the ocean, find the major currents to actively escape predator-rich coastal waters (Scott et al., 2014; Putman & Mansfield, 2015; Putman et al., 2016), and disappear for a period known as the “lost years” (Carr, 1987). At sexual maturity, turtles return to their natal rookery likely using a combination of geomagnetic and olfactory cues (Brothers & Lohmann, 2015; Cameron et al., 2019). Given this strong site fidelity, it is not surprizing that some genomic regions demonstrate patterns of local adaptation (Stiebens et al., 2013).

Overall, despite philopatry, sea turtles have colonized various habitats over evolutionary timescales (Bowen & Karl, 2007). The loggerhead species (*Caretta caretta*) in particularly is widely distributed in tropical and temperate regions, with nesting aggregations ranging from South Africa to Virginia (USA), including the world’s largest rookeries located in Florida (USA, ∼50.000 nests per year) and Masirah Island (Oman; ∼30.000 nests/year, (Baldwin et al., 2003; Bolten & Witherington, 2004; Shamblin et al., 2014). The biogeography of Atlantic loggerheads was hypothetically shaped by geological and climatic events (Bowen & Karl, 2007). The first of these events is the closure of the Isthmus of Panama that separated the Atlantic from the Pacific ∼4.1 M years ago (Duchene et al., 2012; Shamblin et al., 2014). Since then, it has acted as a barrier for species that cannot tolerate freshwater conditions, preventing the movement between the Atlantic and the Pacific Oceans (Bowen et al., 1994; Bowen & Karl, 2007; Zanden et al., 2014). The second major biogeographic event refers to the warm water intrusions that have occurred during interglacial periods around the tip of South Africa, originating from the Agulhas Current (Turney & Jones, 2010). These inflows may have permitted the movement of loggerhead turtles from the Indian Ocean during interglacial periods. Reversely, when the cold Benguela current predominates in the region, low-temperature tolerance may prevent exchange between both ocean basins (Bowen & Karl, 2007).

There are currently two main hypotheses that explain the colonization history of loggerhead rookeries in the Atlantic. On the one hand, it has been proposed that the American rookery, ranging from southern Florida to Northern Carolina, is the oldest rookery in the Atlantic and among the firsts colonized - a conclusion drawn from the high haplotype and nucleotide diversity detected in this aggregation (Reis et al., 2010). On the other hand, Shamblin *et al*. (2014) suggested that the present Brazilian rookeries are the oldest in Atlantic, an hypothesis supported by the basal position of Brazilian mitochondrial DNA (mtDNA) haplotypes in a very large haplotype network (Shamblin et al., 2014). Either way, those events are reflected in the loggerheads’ phylogeny and population structure, which is characterized by the existence of two divergent mtDNA haplogroups as well as genetic differentiation among rookeries within the ocean basin (Bowen et al., 1994; Bowen & Karl, 2007; Shamblin et al., 2014),.

Interestingly, the role of the Eastern Atlantic rookery in a colonization scenario remains to be completely understood (Shamblin et al., 2014). The Eastern Atlantic supports the third largest nesting aggregation of loggerhead turtles in the Archipelago of Cabo Verde (Marco et al., 2012). The Cabo Verde archipelago is located approximately 500km off the Western coast of Africa. It consists of 10 volcanic islands with the oldest ∼20My in the East (Maio) and the youngest aged of ∼8My in the West (Pim et al., 2008). There, turtles lay ∼15.000 nests per year (Marco et al., 2012). The majority of nesting events occurs on the island of Boavista, Maio, Sal and tends to reduce westwards. The existence of two very divergent lineages suggests that at least two independent colonization events occurred (Monzon-Arguello et al., 2010; Stiebens et al., 2013). Similarly, the asymmetric distribution of turtle density in the archipelago calls for the investigation of the directionality of gene flow to better understand the pattern of distribution of genetic diversity. Such knowledge will determine the source and sink nesting groups, facilitating management of this rookery. Nesting density on a smaller geographic scale is also heterogeneous: a beach of 15km length along the south eastern coast of Boavista island supports around >50% of all nesting activity in Cabo Verde (Marco et al., 2012).

In this study, we conducted population genetic analyses on loggerhead rookeries ranging from the large geographic scale of the Atlantic Ocean and Mediterranean Sea, to the regional scale of the Cabo Verde archipelago, and to the local scale of the Island of Boa Vista. We aimed to 1) revisit hypotheses of Atlantic colonization in order to clarify the role of Cabo Verde rookery in the process using hypothesis-driven population genetics modelling, 2) to determine the impact of philopatry on genetic diversity and demographic parameters at various geographical scales from the archipelago to an island level.

## Material and methods

### Global screen for mitochondrial sequences

Sequences of the mitochondrial control region were obtained both from previously published studies as well as by our own data collection for Cabo Verde. The objective was to obtain a robust representation of the rookeries of loggerhead turtles in the Atlantic and the Mediterranean Sea. We retrieved 521 sequences from rookeries in the Mediterranean Sea (Carreras et al., 2007; Clusa et al., 2013; Garofalo et al., 2013; Clusa et al., 2014), 2107 from the USA, which included the whole South Eastern Coast from South Florida to North Carolina (Shamblin et al., 2014) (Monzon-Arguello et al., 2010), 131 for the Brazilian rookery (Shamblin et al., 2014), 175 from Mexico (Shamblin et al., 2014) and 392 others from published studies in Cabo Verde (Monzon-Arguello et al., 2010; Stiebens et al., 2013; Shamblin et al., 2014). In total, 3326 sequences were obtained from published literature (supplementary file S1).

### Field sampling, DNA extraction and sequencing of mitochondrial control region from Cabo Verde

To complement the dataset and improve the resolution at the regional level, field surveys took place in the Cabo Verde archipelago in 2011, 2012 and 2013 during the nesting seasons from June to October. Nine different islands were samples: Boavista, Fogo, Maio, Sal, Santa Luzia, Santiago, Santo Antao, São Nicolau and São Vicente. On the island of Boavista, where the turtle density is the highest (Marco et al., 2012), we sampled on eight different beaches in order to investigate the genetic structure at a local scale. In total, we collected 881 samples from nesting loggerhead females. Samples consisted in removing 3mm piece of non-keratinized tissue from the right front flipper using a single-use disposable scalpel. Samples were immediately stored in ethanol. Turtles were tagged with metal tags and/or pit tags directly after egg deposition to track nesting behaviour and to avoid multiple sampling (Stiebens et al., 2013). In the laboratory, each sample was washed in distilled water for about 20 seconds and cut into smaller pieces. DNA was extracted using the DNeasy® 96 Blood & Tissue Kit (QIAGEN, Hilden, Germany). Elution was conducted in twice 75 μl of AE Buffer. All other steps followed the manufacturer’s protocol.

The long fragment (∼800bp) of control region of the mitochondrial DNA was amplified using the Primers LCM15382 (5’-GCTTAACCCTAAAGCATTGG-3‘) and H950 (5’-GTCTCGGATTTAGGGGTTTG-3‘) (Monzon-Arguello et al., 2010). A 10 μl PCR reaction consisted of 1μl 10x Buffer, 1 μl dNTP’s (10 mM), 0.3 μl MgCl_2_ (5nM), 3.6 μl HPLC water, 0.1 μl Taq Polymerase (Invitek®), 1μl of each primer (5pmol/μl) and 1μl template DNA (∼20μg). The reactions were carried out under the following thermo-cycling conditions: An initial denaturation step of 95°C for 2 minutes, followed by a second cycle that was repeated 40 times with denaturation at 95°C for 30 seconds, annealing at 55°C for 30 seconds and elongation at 72°C for 1 minute. A final elongation step of 7 minutes at 72 °C was carried out. PCR products were cleaned with ExoSAP-IT® following the manufacturer’s protocol. Cycle sequencing reactions were performed with Big Dye® Terminator v3.1 Cycle Sequencing Kit (Applied Biosystems, Darmstadt, Germany). Sequences were obtained from the forward direction (primer LCM15382). Where insufficient fragment lengths were retrieved, sequences from the reverse direction were also obtained and sequences were concatenated into contigs. Sequencing was performed with an ABI 3730 Genetic Analyzer (Applied Biosystems, Darmstadt, Germany). Sequences were assembled in Codon Code Aligner v5.0 (CodonCode Corporation, Dedham, Massachusetts) and ambiguities were corrected by hand. All the amplified mitochondrial sequences were classified accordingly to the standardized nomenclature of the Archie Carr Centre for Sea Turtle Research (http://accstr.ufl.edu). The entire data set was aligned in Muscle v8.3.1 (Edgar, 2004). All unique sequences can be found in supplementary file S1.

### On the large-scale rookeries of the Atlantic basin

#### Genetic diversity and population structure

Haplotype (Hap) and nucleotide diversity (π) indices were computed in Arlequin v3.5.1.3 for each of the major rookeries in the Atlantic and Mediterranean Sea (Excoffier et al., 2005).. Relationship among haplotypes was investigated in a neighbour-joining network using Network v4.6.1.3 (http://www.fluxus-engineering.com), and visualized with PopArt (http://popart.otago.ac.nz). Population structure amongst those major rookeries was investigated using F_ST_ estimates in Arlequin v3.5.1.3 (10.000 permutations). Because the Cabo Verde rookery has still not been properly placed in the broader context of rookeries’ structure, we performed a hierarchic Analysis of Molecular Variance (AMOVA) considering different grouping scenarios, i.e. grouping Cabo Verde with all rookeries separately. The considered rookeries were Brazil, Mediterranean Sea, Mexico and USA. Most likely grouping was identified based on the F_CT_.

#### Testing the ancestry of Atlantic and Mediterranean rookeries

Rookeries’ ancestry and colonization routes within the Atlantic basin and the Mediterranean Sea were explored by comparing the likelihood ratios of model-based inferences. Phylogenetic models were alternatively built with either the Mediterranean, Mexican, Cabo Verdean, USA and the Brazilian rookery as fixed at the root of the tree. The initial tree root height was set to initial split between the genus *Caretta* and the genus *Lepidochelys*, 4.09 million years before present (Duchene et al., 2012). Monophyly was enforced in all rookeries in order to constrain tree topology during the course of MCMC sampling. These analyses were performed in BEAST v.1.8 (Drummond & Rambaut, 2007) and based on haplotype sequences without considering their frequencies. The substitution model was set to HKY as it was found to be the best-fit model of nucleotide substitution chosen through Akaike Information Criteria (AICc) in Mega v6.06 (Tamura et al., 2013) and the mutation rate was set to 3.24*10^-3^ substitutions/site/million year, as estimated for marine turtles (Duchene et al., 2012). The tree prior was set to coalescent and constant size. The MCMC chain length was set to 10^8^ steps. Convergence was inspected in Tracer v1.6 (Rambaut et al., 2014), and models were compared by applying the AICM criteria (1000 bootstraps) (Baele et al., 2012).

#### Testing colonization scenarios within the Atlantic Basin

To complement the Bayesian phylogenies intended to infer the most likely ancestral rookery, we built possible colonization models in order to 1) strengthen phylogenetic conclusions and 2) infer colonization routes within the Atlantic basin. Colonization hypotheses were tested with models that weight the roles of migration, i.e. gene flow, and mutations as sources of genetic novelty within a population. Colonization models were built and compared in the software Migrate-n v.3.6 through Bayesian inference (Beerli & Felsenstein, 2001). Models consisted of different scenarios of rookeries serving as source of migrants, from which only emigration was allowed to occur. In total, we explored 12 possible colonization hypotheses (Fig. S1). Due to the extensive computation power required, we used only unique haplotype data in our preliminary screen. For these models, prior distribution for *gene flow* (M) and *effective population size* (θ) were set as uniform with upper and lower boundaries explored by preliminary tests (θ = 0 - 20, M =0 - 200). Thermodynamic integration with 4 chains with different temperatures (1.0, 1.5, 3.0 and 1000000.0) was performed in order to improve the search for parameter space and allow comparisons of models with Bayes factor. A total of 5×10^5^ steps were recorded in each chain after a burn-in of 10^4^ states. Three independent replicates were performed for each scenario within each run, in order to ensure convergence. A total of 1.5×10^8^ parameters values were visited. Marginal likelihoods were used for model comparisons with Bayes Factor (Kass & Wasserman, 1995).

We further explored colonization hypothesis with approximate Bayesian computation implemented in the software DIYABC v2.1.0 (Cornuet et al., 2014). DIYABC allows the generation of simulated datasets and selection of those closest to the true dataset, and the estimate posterior distribution of specific statistics. The objective was to explore the possibility that the two major mtDNA haplogroups (CC1A and CC2A) may have distinct evolutionary and colonization histories after the split from a common ancestor. The genetic composition of contemporary Atlantic rookeries would therefore reflect several instances of secondary contact between proto-populations composed by individuals carrying haplotypes belonging to major haplogroup CC1 and CC2. DIYABC scenarios were built to test both migrate-n the 3 highest rank models and two others that could not be tested with migrate-n due to increased structure complexity.

In total, we constructed and compared 5 scenarios (Fig, 1). Reference tables were built with 40.000.000 simulated datasets. Runs were optimized to search for the summary statistics with the least distance between simulated and observed dataset. Hence, we used all F_ST_ pairwise comparisons between rookeries from the observed dataset to situate our data with the simulated parameter spaces of the 5 scenarios. Priors were defined as following: uniform population sizes min=100, max=500; uniform branching times (in generations) calibrated for the Last Glacial Maximum (LGM) at t1, considering turtle generation time of 100 years and LGM taking place around 8000-12000ya. Thus, t1 min=80, max=100; t2 min=2000, max=3999; t3 min=4000, max=5999; t4 min=6000, max=8999 and t5 min=9000, max=10000. Mutation rate was also allowed to vary uniformly between 10^-3^ and 10^-7^, with substitution model Kimura-2.

**Figure 1.**
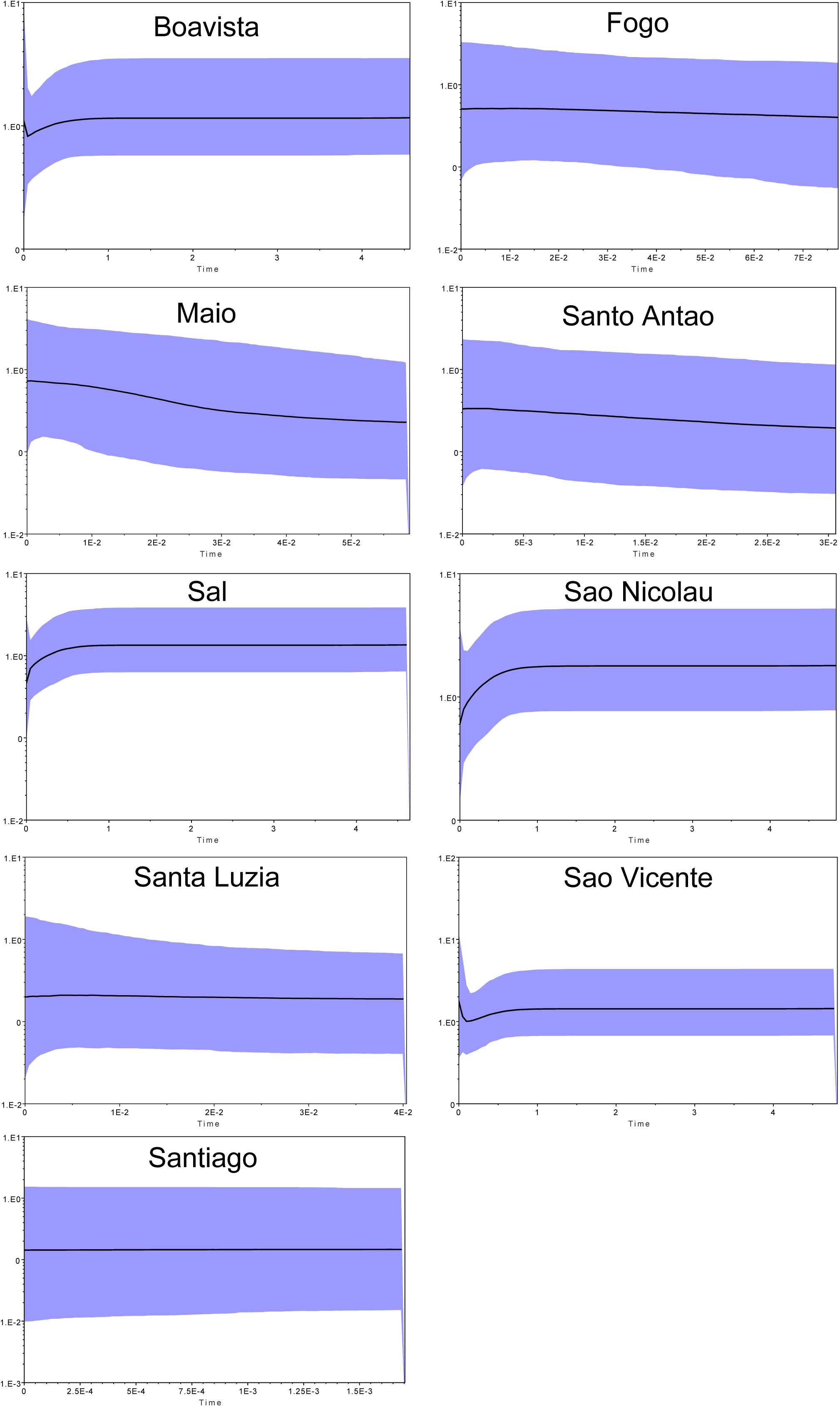
Explored scenarios considering two colonization routes into the Atlantic. The most complex scenarios drawn to test the hypothetical two waves of colonization stemming from two ancestral populations composed by the major haplogroup lineages are depicted. Time points (*t0* to *t5*) of each split are the same across scenarios and are all relative to the initial split that occurred at t=5. Proto populations refer to ancestral populations prior to lineage admixture; *ra* and *1-ra* refer to the ratio of individuals from each ancestral population used to originate the new populations. The pairwise F_ST_ between rookeries estimated from each scenario were then compared to the observed pairwise F_ST_. The scenario best explaining F_ST_ distribution is depicted with a star.

### Regional level of the Cabo Verde archipelago

#### Genetic diversity and structure within the Cabo Verde archipelago

Arlequin v3.5.1.3 was used to calculate the nucleotide and haplotype diversity for each nesting group (i.e. each island), to compute Wright’s fixation index (F_ST_) and perform exact tests for estimation of population differentiation (10.000 permutations). With the exception of calculating genetic indices, and to not influence direct comparisons due to exceptionally high sample size of Boavista, we randomly picked 100 sequences from Boavista and kept them for all downstream analyses. Results were corrected for multiple comparisons using the modified false discovery rate (FDR) method (Benjamini & Yekutieli, 2001), as suggested by Narum (Narum, 2006). Furthermore, we calculated the average F_ST_ for each nesting group in order to investigate whether a geographic pattern of population structure exists across the archipelago.

#### Demographic history and colonization scenarios within the Cabo Verde archipelago

The demographic history of each nesting group was first investigated through moment estimates Tajima’s D (computed with 1000 coalescent simulations), sum of squares deviation (SSD) and a measurement of goodness-of-fit, the raggedness index *r*. All these analyses were performed in Arlequin v3.5.1.3 and DNAsp v5.10(Librado & Rozas, 2009). Then, Bayesian skyline plots were constructed to infer fluctuations of effective population size throughout a temporal scale for each nesting group on each island. These were computed in BEAST v.1.8 (Drummond & Rambaut, 2007). The parameters substitution model and mutation rate were the same as the ones used in the phylogenetic scenarios. The initial tree root height and tree priors were also estimated in these analyses to have another perspective over colonization time for each island. Convergence was inspected in Tracer v1.6. Graphical display of the skylines was constructed in Tracer v1.6.

In order to further investigate the migration along the archipelago’s West-East axis, we calculated the effective number of migrants (ENI) per generation across islands and related it to geographic distance and direction with a linear model (Stiebens et al., 2013). Migration estimates were obtained with migrate-n. For this model, prior distribution for *gene flow* (M) and *effective population size* (θ) were set as uniform with upper and lower boundaries of θ= 0 – 100 and M =0 - 1000. A total of 10^6^ steps were recorded in each chain after a burn-in of 10^4^ states. Two independent replicates were performed for each scenario within each run. A total of 2×10^8^ parameters values were visited. We ran it three times, averaged θand M, and calculated effective migration rates (ENI) with the equation (θ_average_ *M_average_)/2. Geographic distances were calculated from the GPS coordinates of each island. Direction was inferred in relation to the longitudinal position of each island pair. ENIs and geographic distances were log transformed and incorporated in a linear model as response and explanatory variables, respectively, while direction (eastwards or westwards) of gene flow was incorporated as a factor. Statistical analyses were conducted in R v3.2.3 (Team, 2013).

### Within beaches of Boavista

#### Genetic diversity and structure within the island of Boavista

Fine-scale variation in the distribution of genetic diversity was investigated for turtles nesting on the island of Boavista across 7 different beaches. Boavista is the eastern most island of the Archipelago and has an area of 631.1 km^2^. It is the Cabo Verdean Island where the majority of the nesting events takes place (Marco et al., 2012). Diversity indices (haplotype and nucleotide diversities), pairwise F_ST_ comparisons and exact tests of population differentiation amongst nesting beaches were computed in Arlequin v3.5.1.3. Bayesian skylines were performed as mentioned above. Test were corrected for multiple testing.

## Results

Because some sequences retrieved from literature, as well as some obtained in this study, had different lengths, all sequences were trimmed to a consensus length of 674bp. This length does encompass the most polymorphic region of the control region (Monzon-Arguello et al., 2010). In total, we kept 521 sequences from the Mediterranean, 2107 from the USA, 131 from Brazil, 175 from Mexico and 1273 from Cabo Verde. In total, this study includes 4207 different sequences (Supplementary file S1).

### On the large-scale rookeries of the Atlantic basin

#### Diversity and population structure

Indices of genetic diversity revealed that the Mexican loggerhead rookery exhibits the highest haplotype diversity (*Hd*_Mexico_ = 0.770), while the USA rookery holds the highest value of nucleotide diversity (*π*_USA_ = 0.023). The Mediterranean rookery showed the lowest values of haplotype (*Hd*_Mediterranean_ = 0.348) and nucleotide diversity (*π*_Mediterranean_ = 0.001), suggesting this rookery to be the most recently colonized or to be the result of fewer events of colonization. The Cabo Verdean rookery showed one of the highest indices of haplotype diversity (*Hd*_Cabo Verde_ = 0.572) but one of the lowest of nucleotide diversity (*π*_Cabo Verde_ = 0.005) (Table 1). This contrasting reflects the important of placing the Cabo Verde Archipelago in the colonization history of the Atlantic Ocean by loggerhead turtles.

**Table 1.** Diversity indices of major nesting aggregations of the Atlantic Ocean, Abbreviations stand as following: n=number of individual analysed, nHap=number of haplotypes, Hd=haplotype diversity, π=nucleotide diversity.

| *Rookery* | *n* | *nHap* | *Hd* | $\pi$ |
| --- | --- | --- | --- | --- |
| Brazil | 131 | 3 | 0.467 | 0.001 |
| Cabo Verde | 1273 | 20 | 0.572 | 0.005 |
| Mediterranean | 521 | 13 | 0.348 | 0.001 |
| Mexico | 175 | 14 | 0.770 | 0.015 |
| USA | 2107 | 23 | 0.555 | 0.023 |

Not surprisingly given the philopatric nature of loggerhead turtles, pairwise F_ST_ comparisons among global rookeries revealed highly significant differentiation with F_ST_ values ranging from 0.964 (p < 0.001) between the rookeries of Brazil and the Mediterranean Sea to 0.303 (p<0.001) between the rookeries of Cabo Verde and that of the USA (Fig. 2, Table S1). Exact tests of population differentiation showed similar results (Table 2). On average, the USA rookery showed the lowest, though highly significant, level of pairwise differentiation among all rookeries. None of the grouping scenarios tested with hierarchical AMOVA revealed significant F_CT_, suggesting that this approach is not informative enough to infer the overall structure among rookeries.

**Figure 2.**
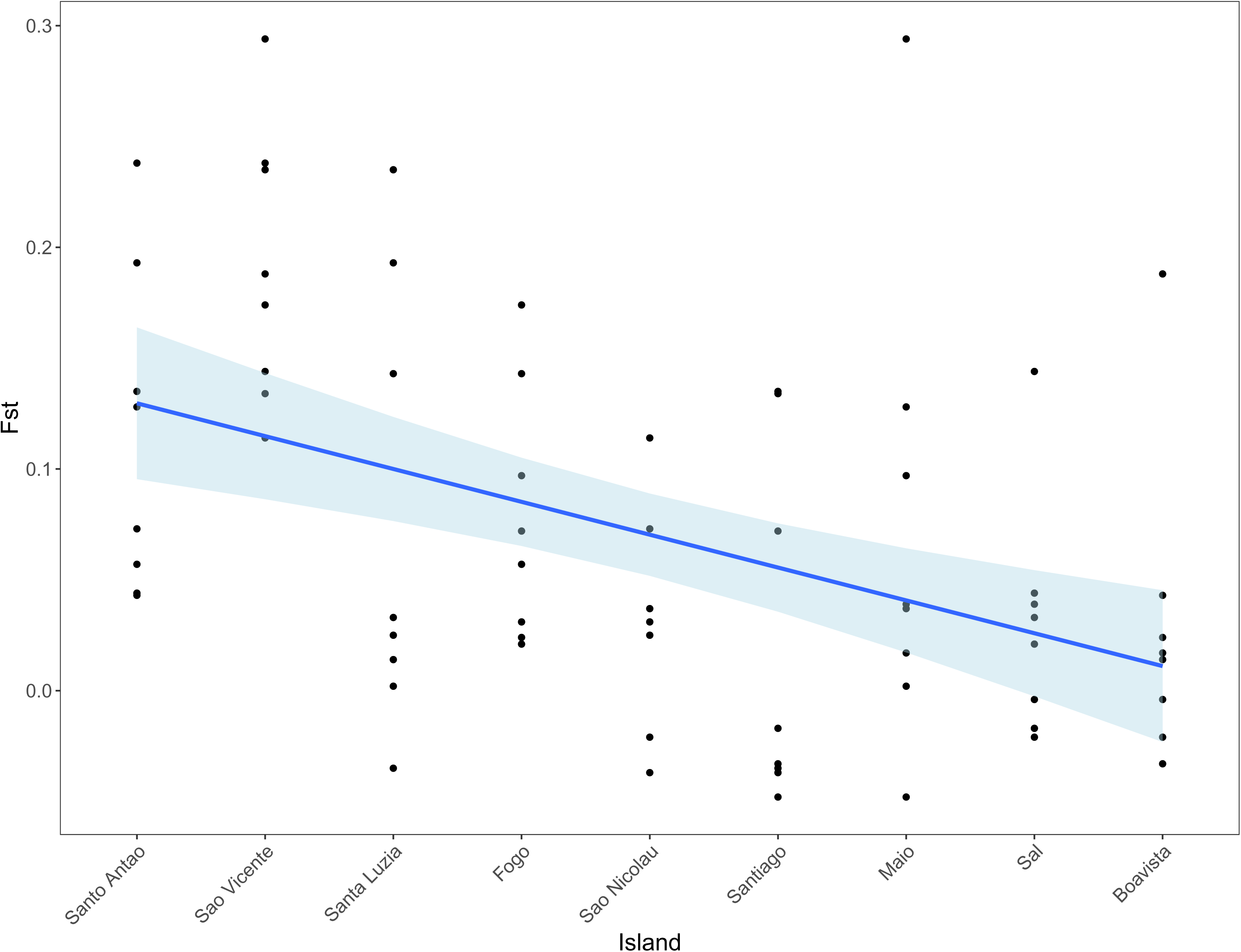
Pairwise F_ST_ distribution across the three geographic scales. Heatmap of the pairwise F_ST_ among (a) major rookeries in the Atlantic basin, (b) islands of the Cabo Verde archipelago, and (c) beaches of Boavista (c). Statistically significant pairwise comparisons are represented with *** for p<0.001 and with ** for 0.001<p<0.01.

**Table 2.** Exact population differentiation tests among rookeries. In bold, significant values for p<0.01.

|  | *Mexico* | *USA* | *MED* | *CV* |
| --- | --- | --- | --- | --- |
| USA | **0.000** |  |  |  |
| MED | **0.000** | **0.000** |  |  |
| CV | **0.000** | **0.000** | **0.000** |  |
| BRA | **0.000** | **0.000** | **0.000** | **0.000** |

#### Ancestry and colonization routes of global rookeries

To Infer ancestry and colonization routes across the Atlantic and into the Mediterranean Sea, we used phylogenetic models, where ancestry and monophyly were enforced sequentially for all rookeries. The comparison of marginal likelihoods of phylogenetic models with fixed ancestry suggested the USA rookery to be the oldest among all the ones analysed (Table S2). Noteworthy, all models showed very similar marginal likelihoods indicating weak statistical differences. Ranking AICM values across models strongly suggest the Mediterranean rookery to be youngest. Comparisons of migrate-n models based on Bayes factors revealed a model with an ancestral Mexico rookery from which turtles colonized all others, as well as a young Mediterranean rookery and a central Cabo Verde rookery acting as a stepping stone towards both sides of the Atlantic (best fit model M4, probability of 0.63, Fig S1, Table S3).

Approximate Bayesian computations revealed scenario S5 to be the most likely (Fig. S3). Scenario S5 suggests Brazil to be the most ancient rookery in the Atlantic, founded by an ancestral haplogroup I population that later colonized the Cabo Verdean rookery before a return towards the area of USA/Mexico. This S5 scenario further implies that the Mediterranean Sea was the last rookery to be founded but only by haplogroup II, whose individuals dispersed then to USA/Mexico and from there to Cabo Verde archipelago. Model inferences considering a lineage split prior to colonization and dispersal were never been attempted, and here we show that doing it allows the explore the likelihood of theorized colonization scenarios.

### Regional level of the Cabo Verde archipelago

With Cabo Verde appearing as a central rookery acting as a stepping stone along the colonization of the Atlantic basin, we investigated the demographics, diversity and population structure of this rookery. For the following analyses, we used 1273 mtDNA sequences of nesting female turtles. Twenty-two different haplotypes were detected, among them two haplotypes (UH5 and UH13, Supplementary File S1) that were found in previous study (Stiebens et al., 2013b) but not yet described in Genbank.

#### Demographic history of the archipelago

Because demographic history may alter the understanding of the colonization patterns, we investigated the variation in various demographic indices for each nesting group (i.e. island) within the rookery. Results revealed two distinct patterns. The first one describes a possible population expansion in most of the northern set of islands of Sal, Santo Antao, Sao Nicolau, and Boavista. Together, with Fogo, all Tajima’s were negative, non-significant SSD and raggedness indices (Table 3). However, only Sao Nicolau exhibited a significant Tajima’s D. The other nesting groups rather experienced a constant population size or a decline as seen in Sao Vicente, Santiago, Santa Luzia and Maio islands (Table 3), and amongst those, only Sao Vicente showed a significant Tajima D.

**Table 3.** Diversity indices for the islands of Cabo Verde. Abbreviations stand as following: n=number of individual analysed, nHap=number of haplotypes, Hd=haplotype diversity, π=nucleotide diversity, SSD=sum of squared differences from mismatch distribution, r=raggedness index. In bold, significant values for p<0.05.

| *Island* | *n* | *nHap* | *Hd* | $\pi$ | *Tajima's D* | *SSD* | *r* |
| --- | --- | --- | --- | --- | --- | --- | --- |
| Boavista (BV) | 831 | 16 | 0.564 | 0.004 | -1.486 | 0.067 | 0.269 |
| Fogo (FG) | 26 | 4 | 0.640 | 0.001 | -0.390 | 0.004 | 0.089 |
| Maio (MAI) | 101 | 8 | 0.566 | 0.001 | 0.074 | 0.061 | 0.255 |
| Sal (SI) | 100 | 8 | 0.582 | 0.007 | -0.964 | 0.027 | 0.118 |
| Sao Nicolau (SN) | 24 | 6 | 0.681 | 0.006 | **-2.222** | 0.009 | 0.041 |
| Sao Vicente (SV) | 65 | 10 | 0.662 | 0.020 | **2.189** | 0.112 | 0.171 |
| Santa Luzia (SU) | 55 | 3 | 0.560 | 0.002 | 0.879 | 0.039 | 0.147 |
| Santiago (ST) | 11 | 3 | 0.564 | 0.001 | 1.176 | 0.053 | 0.251 |
| Santo Antao (SA) | 60 | 5 | 0.438 | 0.001 | -0.760 | 0.000 | 0.113 |

Reconstructing the effective population size estimates from Bayesian skyline plots revealed that Sao Vicente, Boavista, Sao Nicolau and Sal showed a recent decline in effective population size followed by a possible recovery close to present time (Fig. 3). A possible demographic recovery is thus in line with negative Tajima D values. Nesting populations on the islands of Fogo, Maio, Santa Luzia, Santiago and Santo Antao instead show a relatively stable effective population size instead, where only in Maio and Santa Luzia did Tajima’s D agreed with Bayesian skylines. Notably and despite all runs attained convergence, the observation that high probability density interval widens close to the present implies uncertainty regarding the current dynamics of each island’s demography.

**Figure 3.**
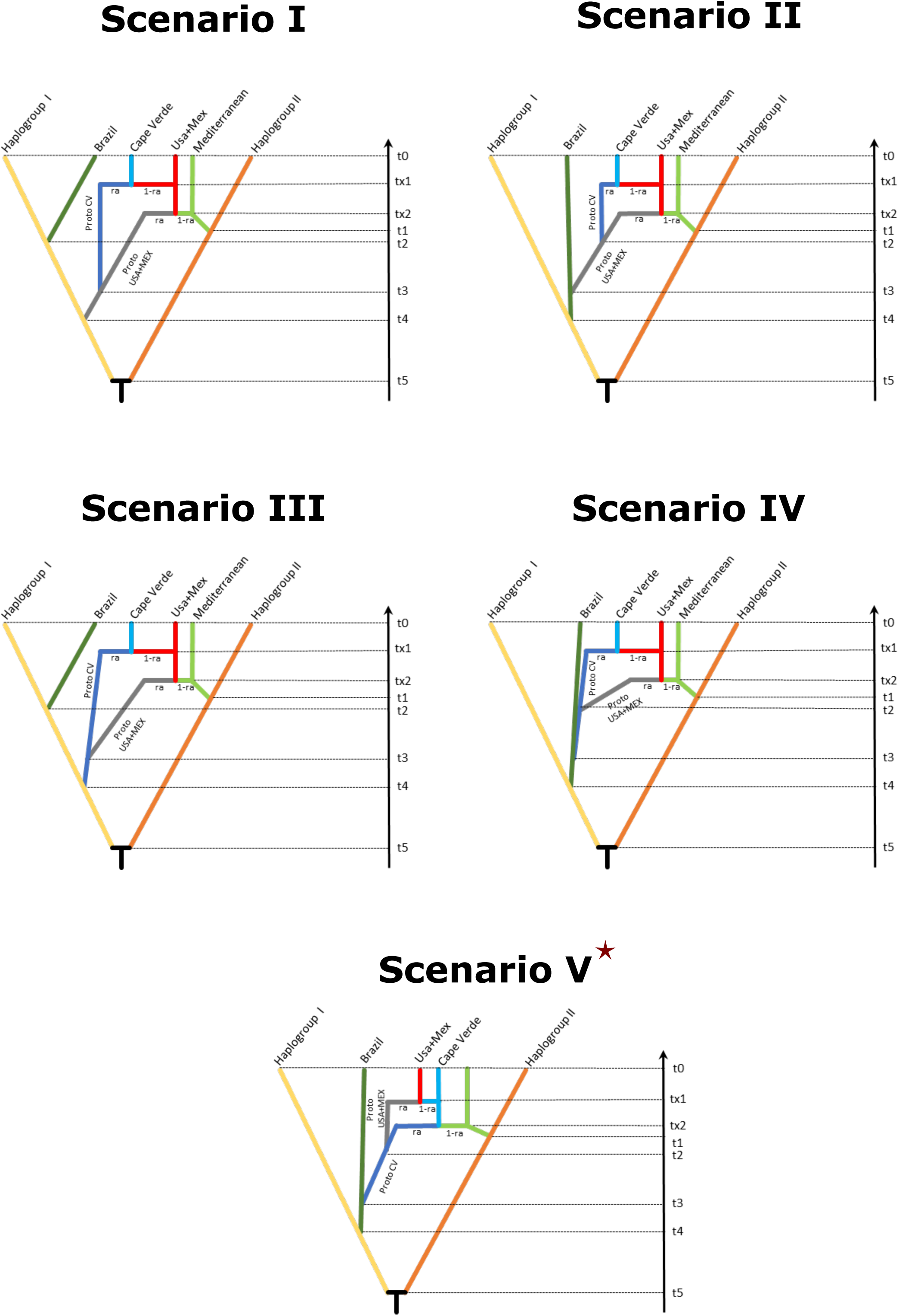
Bayesian Skyline Plots (BSPs) for the nine islands of Cabo Verde. Effective population size across evolutionary timescales for each island. Y-axis is an estimate of the product of *Ne* * mutation rate (µ) per generation, while the x-axis represents a relative temporal scale. Note that the differences in x-axis scale across graphs reflect the putative age of the genealogy. The black like represents the mean *Ne* and the blue shading the 95% high probability density interval. There is an inflection points for Boavista and Sao Vicente populations, suggestive of an increase after a population decline. The islands of Sal and Sao Nicolau present a declining trend and the aggregations on the remaining islands suggest stability of effective population sizes in the respective time frames.

#### Diversity and structure of each island

The most frequent haplotypes were CCA1.3 (n=766) and CCA17.1 (n=294) that belong to the oldest Haplotype I lineage. As expected, haplotype and nucleotide diversities show less variation at the regional level than at the global level, however they remain high and diverse (Table 3). Indices of genetic diversity split for nesting islands showed that turtles sampled in Sao Nicolau harbours the highest haplotype diversity (*Hd*_SaoNicolau_ = 0.681) while those from Sao Vicente showed the highest nucleotide diversity (π_SaoVicente_ = 0.020). Both islands belong to the northern area of the archipelago. The lowest values of haplotype diversity were observed in turtles from Santo Antao (*Hd*_SantoAntao_ = 0.438) and the lowest nucleotide diversities were detected in turtles from four islands: Santo Antao, Santiago, Fogo and Maio (π = 0.001) (Table 3). Noteworthy, the last three islands are adjacent in the South of the archipelago.

Pairwise F_ST_ comparisons among islands resulted in twelve statistically significant comparisons after correction for multiple comparisons (Fig. 2b). The genetic composition of Sao Vicente Island produced the highest F_ST_ values among island pairs, with statistically significant pairwise F_ST_ ranging from 0.174 with Fogo (p = 0.005) to 0.294 with Maio (p < 0.001) (Table S4). Exact tests of population differentiation showed seventeen significant results, mostly consistent with pairwise F_ST_ particularly those involving the islands of Sao Vicente, Fogo and Sal (Table 4). F_ST_ did not correlate with geographic distance (Mantel test: r = -0.138 p=0.820), suggesting an absence of isolation by distance.

**Table 4.** Exact population differentiation tests among islands of the Cabo Verde Archipelago. In bold, significant values for p<0.05.

|  | *Boavista* | *Sal* | *Sao Vicente* | *Sao Nicolau* | *Fogo* | *Maio* | *Santa Luzia* | *Santo Antao* |
| --- | --- | --- | --- | --- | --- | --- | --- | --- |
| Sal | 0.140 |  |  |  |  |  |  |  |
| Sao Vicente | 0.000 | **0.000** |  |  |  |  |  |  |
| Sao Nicolau | 0.039 | 0.195 | 0.076 |  |  |  |  |  |
| Fogo | **0.001** | **0.007** | **0.000** | 0.098 |  |  |  |  |
| Maio | 0.335 | 0.020 | **0.000** | **0.006** | **0.003** |  |  |  |
| Santa Luzia | 0.151 | 0.029 | **0.000** | **0.006** | **0.000** | 0.100 |  |  |
| Santo Antao | **0.000** | **0.000** | **0.000** | **0.009** | **0.000** | **0.000** | **0.000** |  |
| Santiago | 0.933 | 0.764 | 0.237 | 0.623 | 0.326 | 0.777 | 0.747 | 0.015 |

Interestingly, the average pairwise F_ST_ was shown to significantly vary across islands (ANOVA: F =4.367, p<0.001), increasing in a linear fashion from East to West (R^2^=0.275, p<0.001, Fig. 4). To further describe the East-West pattern, we estimated the number of migrants amongst islands and the direction of gene flow. We found a significant interaction between the distance among islands and the direction of gene flow: the average number of migrants decreases eastwards (t_direction West_ =-1.281, p =0.047) and increases westwards (t_direction West_ =0.719, p =0.015). This result demonstrates that the large nesting groups in the east mainly serve as source of migrants in the smaller nesting groups over evolutionary times (Fig. S3).

**Figure 4.**
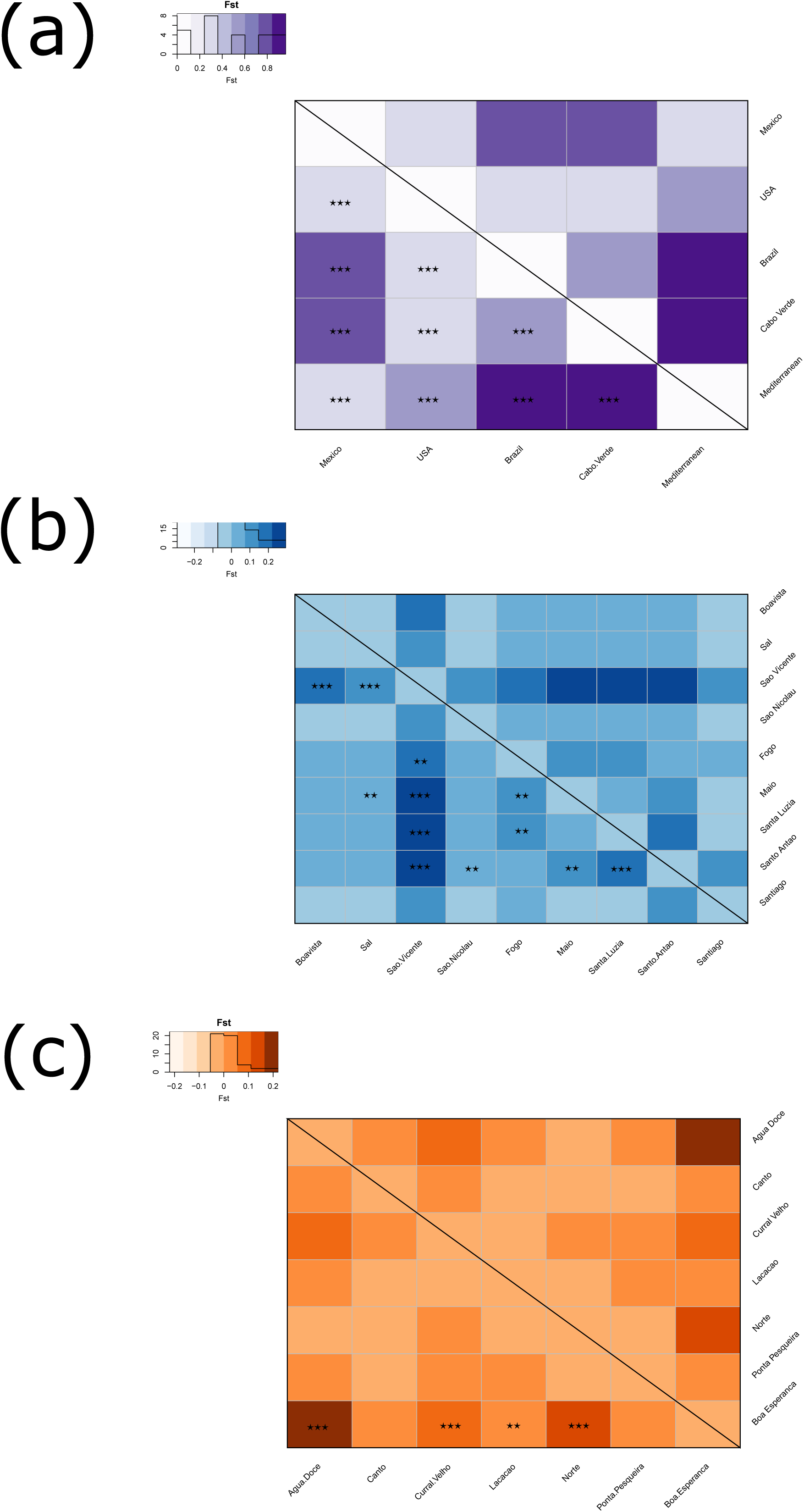
Distribution of average pairwise F_ST_ among the islands of the archipelago. Linear regression of average pairwise F_ST_ per island along the East to West axis of the archipelago following the equation *F_ST_ ∼ Island*. A significant increase in average pairwise F_ST_ can be observed from East to West (R^2^=0.275, p<0.001).

### Within beaches of Boavista

The island of Boavista supports the largest nesting group in the Eastern Atlantic (Marco et al., 2012). We used the largest dataset collected from specific beaches in the island (N=626) to investigate the genetic diversity and population structure at the local scale (Table 1C). We found that the beach of *Agua Doce* showed the highest haplotype and nucleotide diversity (*Hd*_AguaDoce_ = 0.752, *π*_AguaDoce_ = 0.009), while Boa Esperança exhibited the lowest values in both indices (*Hd*_BoaEsperança_ = 0.331, *π*_BoaEsperança_ = 0.001). Interestingly, those beaches are adjacent to each other. Furthermore, haplotype diversity observed at the local scale was comparable to that of the regional level but reduced of about half for the nucleotide diversity (Table 5).

**Table 5.**
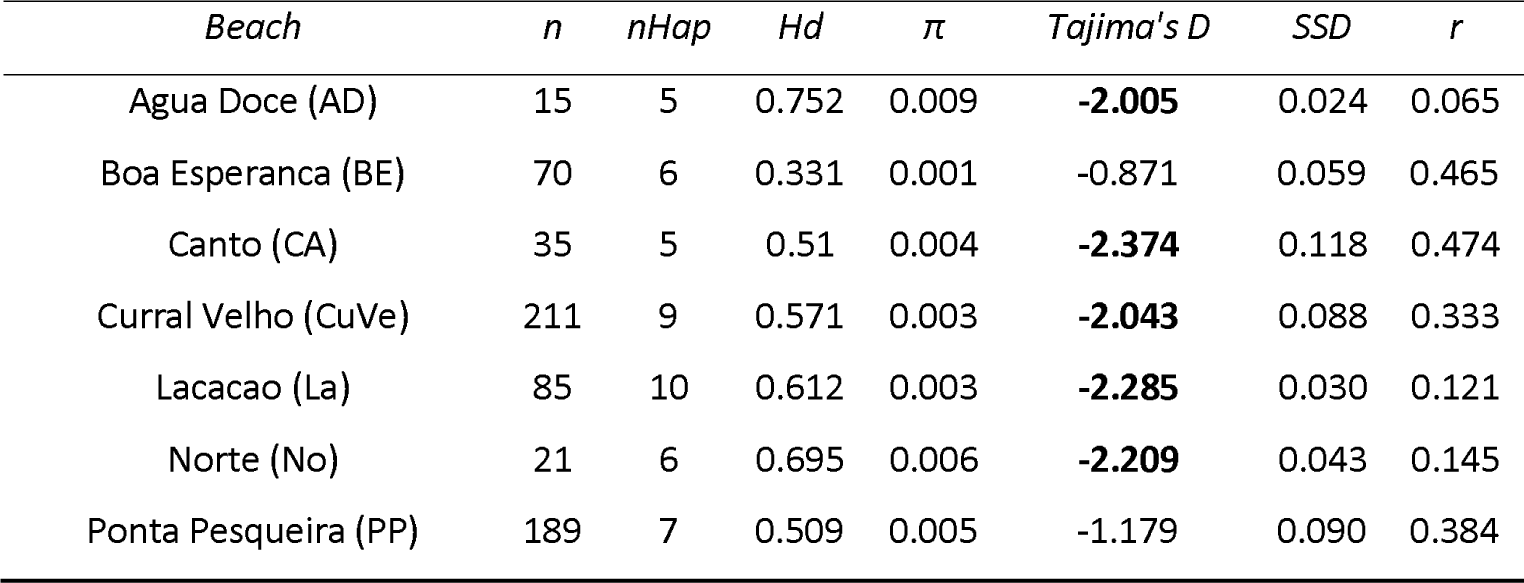
Diversity indices for beach in Boavista. Abbreviations stand as following: n=number of individual analysed, nHap=number of haplotypes, Hd=haplotype diversity, π=nucleotide diversity, SSD=sum of squared differences from mismatch distribution, r=raggedness index. In bold, significant values for p<0.05.

| *Beach* | *n* | *nHap* | *Hd* | $\pi$ | *Tajima's D* | *SSD* | *r* |
| --- | --- | --- | --- | --- | --- | --- | --- |
| Agua Doce (AD) | 15 | 5 | 0.752 | 0.009 | **-2.005** | 0.024 | 0.065 |
| Boa Esperanca (BE) | 70 | 6 | 0.331 | 0.001 | -0.871 | 0.059 | 0.465 |
| Canto (CA) | 35 | 5 | 0.51 | 0.004 | **-2.374** | 0.118 | 0.474 |
| Curral Velho (CuVe) | 211 | 9 | 0.571 | 0.003 | **-2.043** | 0.088 | 0.333 |
| Lacacao (La) | 85 | 10 | 0.612 | 0.003 | **-2.285** | 0.030 | 0.121 |
| Norte (No) | 21 | 6 | 0.695 | 0.006 | **-2.209** | 0.043 | 0.145 |
| Ponta Pesqueira (PP) | 189 | 7 | 0.509 | 0.005 | -1.179 | 0.090 | 0.384 |

**Table 6.** Exact population differentiation tests among the beaches in Boavista. In bold, significant values for p<0.05.

|  | *Agua* |  | *Curral* |  |  | *Ponta* |
| --- | --- | --- | --- | --- | --- | --- |
|  | *Doce* | *Canto* | *Velho* | *Lacacao* | *Norte* | *pesqueira* |
| Canto | 0.028 |  |  |  |  |  |
| Curral Velho | 0.035 | 0.169 |  |  |  |  |
| Lacacao | 0.283 | 0.350 | 0.286 |  |  |  |
| Norte | 0.763 | 0.178 | 0.023 | 0.140 |  |  |
| Ponta |  |  |  |  |  |  |
| pesqueira | **0.002** | 0.085 | 0.023 | **0.001** | 0.011 |  |
| Boa Esperanca | **0.000** | 0.117 | **0.000** | **0.001** | **0.000** | **0.000** |

Surprisingly, pairwise comparisons were found significant at this local scale. All significant tests included the beach of Boa Esperança (Fig. 2c): Boa Esperança and Agua Doce (F_ST_ = 0.220, p = 0.00), Boa Esperança and Curral Velho (F_ST_ = 0.062, p = 0.00), Boa Esperança and Lacacao (F_ST_ = 0.051, p = 0.002) and Boa Esperança and Norte (F_ST_ = 0.114, p = 0.000) (Table S5). Given that we sampled 70 turtles from Boa Esperança, those significant results are unlikely to stem from small sample size.

All beaches of Boavista revealed a pattern of population expansion as suggested by negative Tajima’s D and non-significant SSD and raggedness similar to the apparent scenario of population expansion detected in the above analyses for all the islands (Table 5). Bayesian skylines, however, revealed that Norte, Canto and Agua Doce might have experienced a short decline, while Boa Esperanca experienced a short expansion, when groups of turtles nesting on the other beaches showed no change in population size (Fig. S4).

## Discussion

To effectively manage endangered species, it is crucial to understand the distribution of genetic diversity. In this study, we evaluated the genetic diversity of loggerhead sea turtle populations, known for their natal philopatric behaviours. Philopatry affects the distribution of genetic diversity since it reduces gene flow among populations. Nonetheless, the loggerhead turtle has colonized the entire Atlantic Ocean and the routes underlying this colonization are still elusive. Here, we show support for the Brazilian rookery to be the most ancient rookery in the Atlantic but support the idea that the colonization of the entire ocean basin occurred via two waves: one from the Pacific and other from the Indian. Each colonization event corresponds to that of a major haplotype group, both being now distributed across most of the Atlantic rookeries. Furthermore, we show that Cabo Verde rookery has played a key role in the colonization process, acting as a stepping stone facilitating the establishment of other rookeries on either side of the Atlantic. Focusing on Cabo Verde, we also show that the island supporting the largest nesting density is not necessarily that with the highest diversity indices. This is likely the result of an asymmetric functioning of the different nesting groups of the archipelago, with the Eastern islands acting as sources and the edge of the distribution mostly acting as sink. The eastern islands are those where turtle density is the highest and we find that because of strong philopatry, ∼10kms accuracy, exchanges among islands are relatively rare.

### Revisiting the colonization of the Atlantic: a new perspective reconciles old hypothesis

The existence of two highly divergent mitochondrial haplogroups that shape the phylogeographic patterns of the different ocean basins underlies the loggerhead’s global biogeography (Bowen et al., 1994). The Atlantic being the youngest ocean basin, the unequal distribution of mtDNA haplogroups among rookeries suggests different colonization waves, as those lineages stem from the Indo-Pacific area (Shamblin et al., 2014). Therefore, to understand the colonization and posterior formation of the Atlantic rookeries, we considered those haplogroups as independent ancestral populations rather than forming one joint source of colonization. Modelling the temporal succession of haplogroup dispersal into and within the Atlantic with ABC reconciles current colonization hypotheses under a single scenario. More specifically, the most parsimonious succession of events suggest that the Brazilian rookery was colonized first by individuals carrying haplotypes of major haplogroup I – probably a leakage from the Pacific into the Atlantic while the Central American Seaway was present. This was followed by the colonization of the Cabo Verdean archipelago and then the North and Central American rookeries of USA and Mexico. The first stage of Atlantic colonization is line with line with Shamblin et al (2014)’s hypothesis, as the Brazilian rookery is the oldest of the Atlantic and composed only by the turtles carrying haplotypes of group I. The colonization from Cabo Verde to northern Americas can be explained by the fact that warm saline waters were gradually introduced to northern latitudes as the Central American Seaway shallowed until total closure (Haug & Tiedemann, 1998). As loggerhead turtles require sand temperatures of at least 25° for successful incubation (Bowen & Karl, 2007), North American habitats might not have been suitable for nesting until after the closure of the isthmus of Panama.

On a more recent timescale, individuals carrying haplogroup II colonized the Mediterranean Sea. Likely after the last glaciations, migration occurred to Cabo Verde, where haplogroup II individuals established on already formed colonies, further proceeding to populate USA/Mexico. The pathway here presented thus supports Shamblin’s et al (2014) hypothesis, specifically, the retention in the Indian ocean and subsequent entry via South Africa into the Atlantic (Shamblin et al., 2014). The major haplogroup II pathway favoured by the scenario modelling supports the nearly consensual hypothesis that Mediterranean Sea supports the youngest rookeries with access on the Atlantic Ocean (Clusa et al., 2014).

The biogeographic reconstruction of colonization pathways that assumes two distinct haplogroups as distinct ancestral populations clarifies key elements of the dispersal of loggerheads in the Atlantic. First, it rejects Central and North American rookeries as the oldest in the Atlantic. Notably, our summary statistics such as diversity indices and F_ST_, as well as testing colonization hypothesis without segregating into haplogroups also suggested USA and/or Mexico as the oldest rookery. The apparent paradox can be explained by the physical convergence of the two divergent lineages in two waves of colonization in the area. The admixture of lineages from multiple genetic backgrounds is well known to increase diversity at sink populations (Roman & Darling, 2007). Thus, by being the last rookery to receive individuals carrying the highly diverse haplogroup I, USA/Mexico hold a signature that could be interpreted as that of an ancestral rookery, as observed by Reis et al (2010). Overall, the Atlantic can be understood as a secondary contact zone of major haplogroups that have accumulated divergence in allopatry. While we cannot exclude demographic effects that could have erased presence of genetic variants at local scale, one can parsimoniously assume that once admixed rookeries became established those would affect lineages equally.

### Colonization history, structure and demography of the Cabo Verdean rookery

Regionally, the Cabo Verdean rookery shows both high haplotype and nucleotide diversities with the presence of both mtDNA lineages. Particularly, the island of Sao Vicente supports a nesting group with nucleotide diversity at least 3 times higher than that of other nesting groups of the archipelago. This is mainly linked to the presence of the most divergent lineage being frequent on this island. The presence of this haplotype can be traced mostly to one specific beach called Lazareto on the North-West of the island representing ∼70% of the individuals. The presence of this Indo-Pacific haplotype, also present in the Mediterranean Sea- and interpreted as evidence for a second wave of colonization (Stiebens et al., 2013), is further reinforced by scenario simulations of the Atlantic colonization.

From a structure point of view, Cabo Verde was considered a single nesting population (Monzon-Arguello et al., 2010) and management unit (Wallace et al., 2011; Shamblin et al., 2014), while early signs of differentiations have been detected (Stiebens et al., 2013). The analyses of the more extensive dataset in this study confirms the highly significant genetic differentiation congruent with both F_ST_ and exact tests. This structure arises clearly from the island of Sao Vicente and the frequent divergent haplotype but not only. Indeed, a clear geographic pattern is visible where the islands from the western range of the archipelago show increased average differentiation compared to those of the eastern range. Noteworthy, signatures on mtDNA stem from female-mediated gene flow. Because bi-parentally inherited markers show an increase gene flow eastwards linked to opportunistic mating from Western males encountering more females on the east (Stiebens et al., 2013), our results suggest that the densely-populated easternmost islands of Sal, Boavista and Maio are extremely important in maintaining the functioning of the rookery, acting as evolutionary source populations. Here, we also reveal another geographic pattern, where gene flow of nesting females has propagated from the centre of the distribution to the edge. This pattern might be the ancient signature of colonization within the Archipelago: after the evolution of site fidelity to the firstly colonized islands, populations were established in all others through sporadic nesting events from the long-distance migrants, as it has recently been described to occur in Mediterranean rookeries (Carreras et al., 2018). Altogether, the observed genetic structure is the likely result of a strong female philopatric behavior as often observed in loggerhead turtles (FitzSimmons et al., 1997; Lee et al., 2007).

From a demographic perspective, estimates of effective population size suggest that after a small decline, the largest nesting groups are expanding. This is particularly the case for an East-West corridor entailing Boavista to Sao Vicente islands. While dating the decline is complex, the Cabo Verde rookery may hold the signature of intense poaching which has significantly impacted the census population size (Marco et al., 2008). This signature does not necessarily represent the recent poaching peaks of the 1990s and 2000s. Instead, it may be the footprint of poaching activity, known since the human occupation of the archipelago in the 15^th^ century (Loureiro & Torrão, 2008). However, demographic estimates should be interpreted with caution: the wide high probability density intervals observed for Bayesian skylines and non-significant Tajima D does not support conclusive evidence. Thus, conservation driven actions should be maintained in the Archipelago as well as monitoring of loggerheads census population sizes.

### Philopatry at a fine geographic scale: structure and demography of the island of Boavista

At the local scale, our results show that turtles nesting on one beach of the island of Boavista, Boa Esperança, harbour the lowest nucleotide diversity. This beach is situated about 12kms away from Ponta Ervatao & Ponta Cosme, the beaches with the highest nesting density in Cabo Verde and in the eastern Atlantic (>2500 nest per year, (Marco et al., 2012)). Reduced genetic diversity is likely the result of genetic isolation as demonstrated by the consistent significant differentiation of Boa Esperança with the surrounded beaches. Given that the sample size from this beach is ∼70, this significant differentiation is unlikely to arise from random fluctuation linked to a small sample size. The parsimonious explanation is a high accuracy of philopatric behaviour on Boavista which can be detected away from the centre of the nesting activity, on the other side of the island, as the result of leakage of nesting behaviours.

## Conclusion and future directions

Our results offer a fresh perspective on the biogeography of the loggerhead sea turtle. Understanding the Atlantic as a secondary contact zone for two highly divergent lineages showed that ongoing hypothesis can be merged into a single colonization and dispersal scenario. The expansion pattern exposed allows to investigate major climatic and geological changes that shaped the pathways of colonization - such as the impact of the gradual closure of the isthmus of Panama in opening suitable nesting habitats at Northern latitudes. Our work also suggests that interpreting ancestry from genetic diversity estimates should be made with caution. As we have shown here, the presence of divergent lineages does not necessarily imply ancestry. Hypothesis testing with population genetic modelling may help to clarify conflicting views on the global biogeography.

Despite extensive migration, significant degrees of population differentiation spanned all the analysed geographic levels. Therefore, we have demonstrated that turtles are capable of extreme site fidelity and forming independent nesting groups at highly localized geographic scale. We identified two mechanisms. At the small scale, leakage away from the main nesting sites results in expansion. This mechanism yet relies on large population size, which may be problematic for endangered species. The second mechanism of expansion relates to major climatic/geological events which requires the persistence of the species on a long evolutionary time.

## Acknowledgments

The authors would like to thank the team members and volunteers of BiosferaI, Caretta Caretta, Fondation Biodiversity Maio, National Institute for the Development of Fisheries (INDP), Projecto Vito Fogo, Projecto Vito Porto Novo, Project Biodiversity and Turtle Foundation who assisted with field work. The study was supported by a National Geographic grant (GEFNE69-13) and Queen Mary University Funds allocated to CE. MBS is currently supported by the FCT strategic project UID/MAR/04292/2013 granted to MARE. All applicable institutional and/or national guidelines for the care and use of animal were followed. The work was performed under DGA legislation of Cabo Verde (authorizations DGA 30/13, 040/GP/INDP/14).

## Data accessibility statement

All sequences obtained specifically for this study will be deposited in NCBI’s nucleotide database.

## Author contributions

CE, JDK and MBS designed the study. CSM, RT, TAA, MRS, DPL, DJ, PLJ, DH, CSJK, SVA and CE conducted field work in Cabo Verde. MBS and JDK performed the analysis with CE guidance. MBS drafted the manuscript with CE and JDK. All authors approved the final version of the manuscript.

---

**Figure S1 – Global colonization models** Hypothesized models of Atlantic colonization tested with migrate-n. **BRA**: Brazilian rookeries, **CV**: Cabo Verdean rookeries, **MEX**: Mexican rookeries, **USA**: North American rookeries; **MED**: Mediterranean rookeries. All biologically plausible combinations of unidirectional migration were tested.

**Figure S2 – Logistic regression comparison of scenarios modelled in diyABC** Scenario comparison results of **hypothesized** two colonization routes into the Atlantic. In the x-axis is shown the number of simulated scenarios (in batches of 1000) whose mean pairwise F_ST_ is more approximate to that obtained with the observed dataset. The y-axis is the corresponded regression coefficient for sequential batches of 1000 points.

**Figure S3 – Relationship between migration estimates and geographic distance for each direction** X-axis shows the log-scaled geographic distance and the y-axis shows the effective migration rate for eastward (E) and westward (W) migration.

**Figure S4 – Bayesian Skyline Plots of Bovista’s beaches** Effective population size across evolutionary timescales for each island. Y-axis is an estimate of the product of *Ne* * mutation rate (µ) per generation, while the x-axis represents a relative temporal scale. Note that the differences in x-axis scale across graphs reflect the putative age of the genealogy. The black like represents the mean *Ne* and the blue shading the 95% high probability density interval.

## Notes

#### Summary of Updates

Update author list and newer version.

