## Supplementary figures and images for "Distribution of genetic diversity reveals colonization and philopatry of the loggerhead sea turtles across geographic scales"

### Fig. S1

#### Global and Global with frequency data

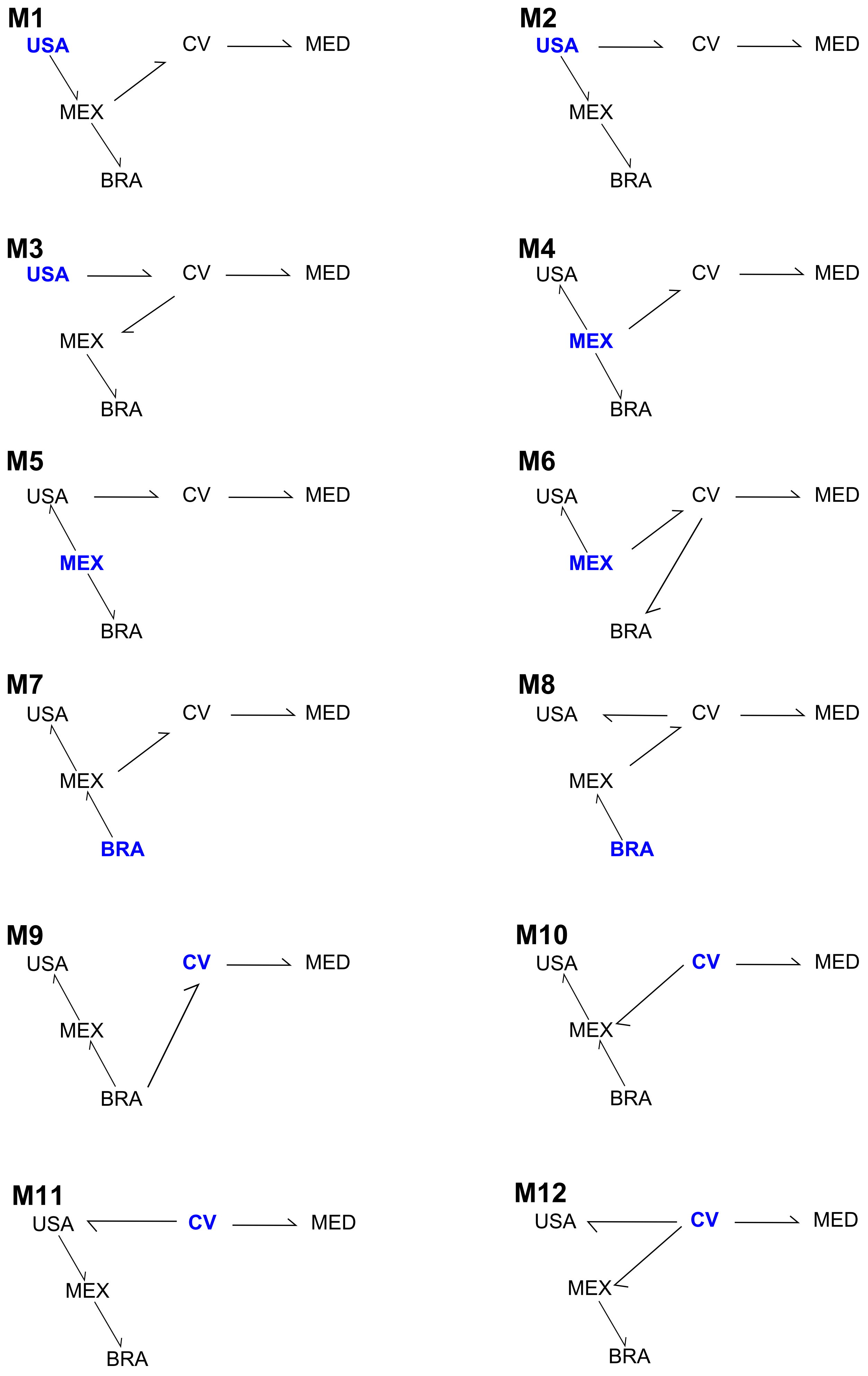

### Fig. S2

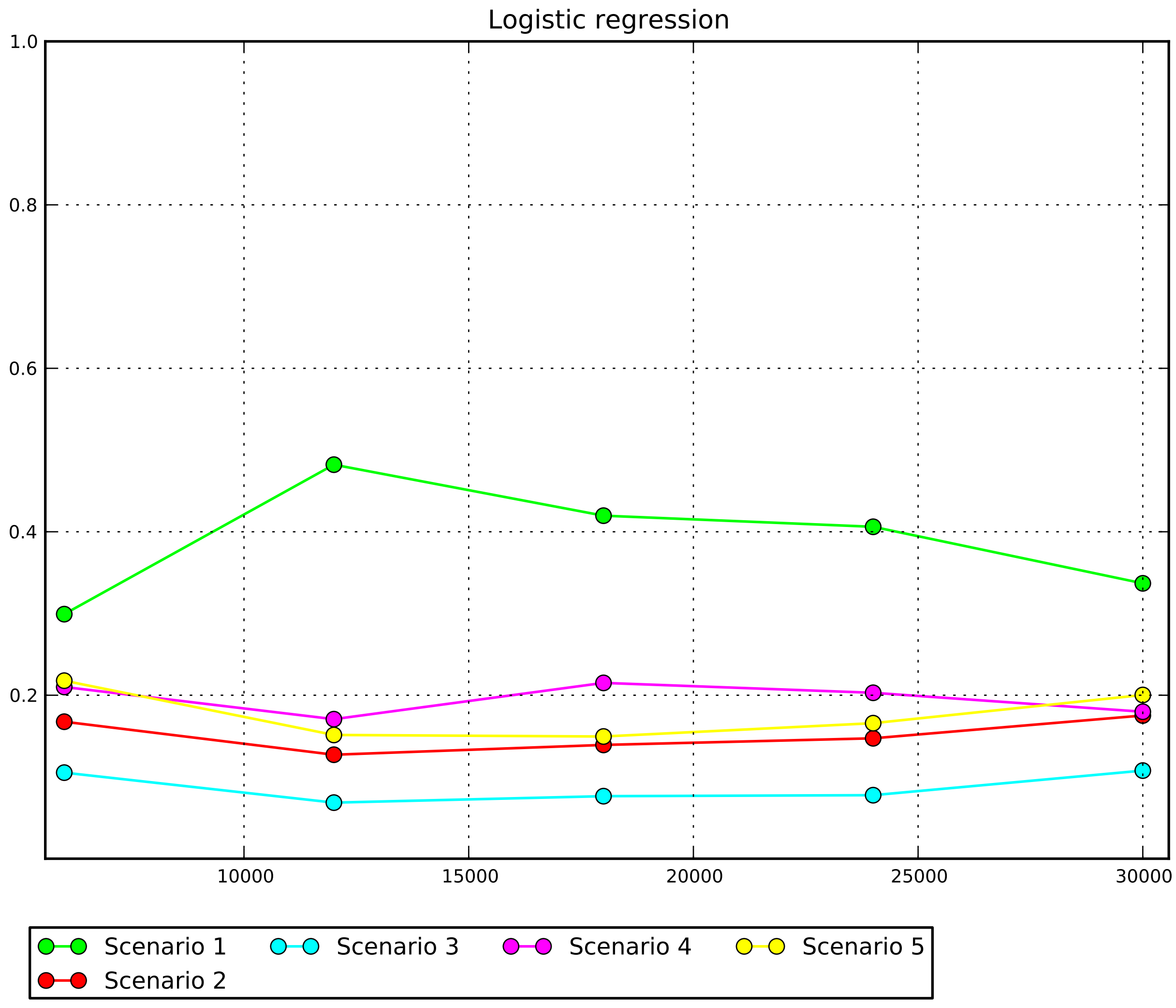

### Fig. S3

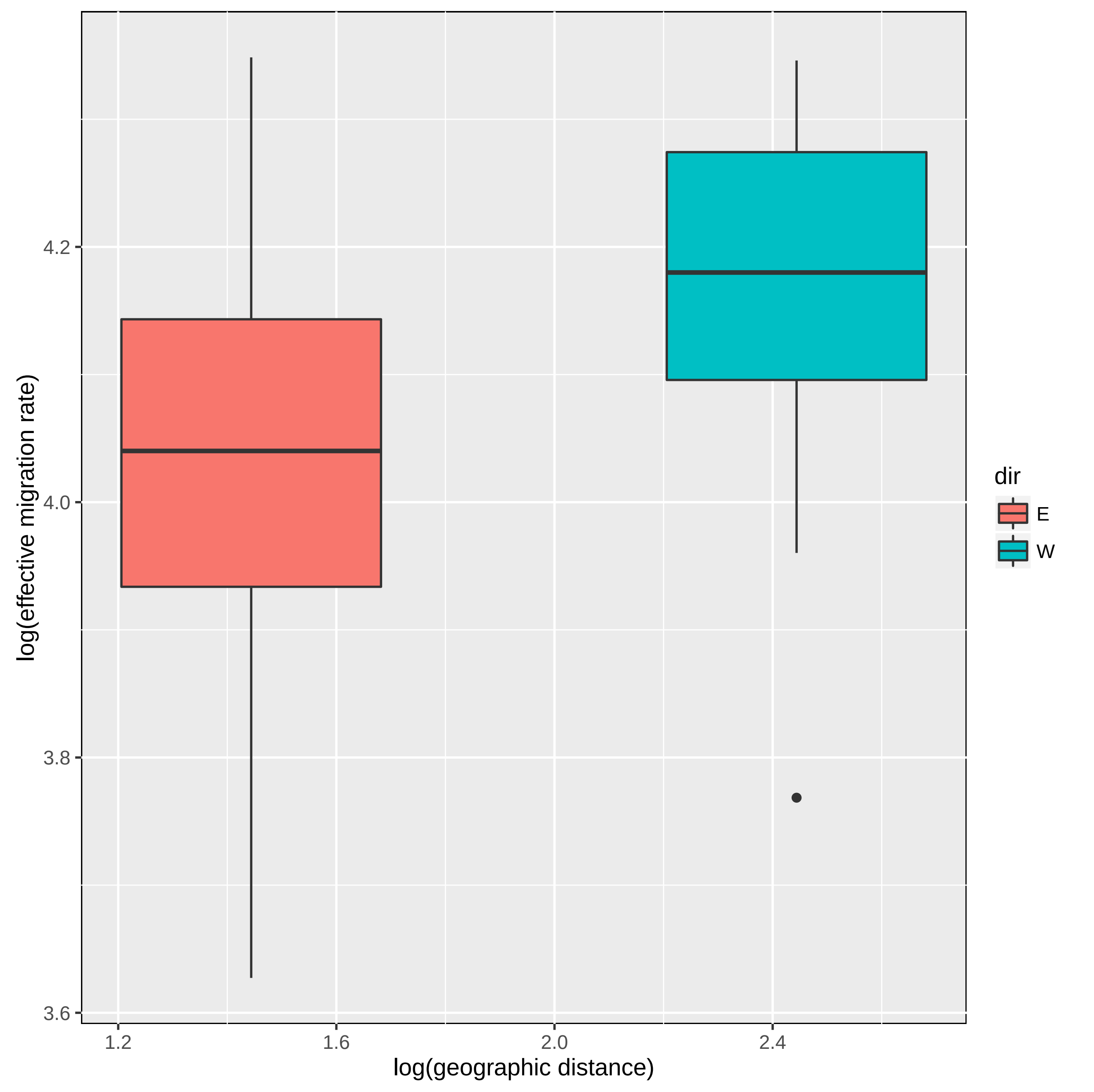

### Fig. S4

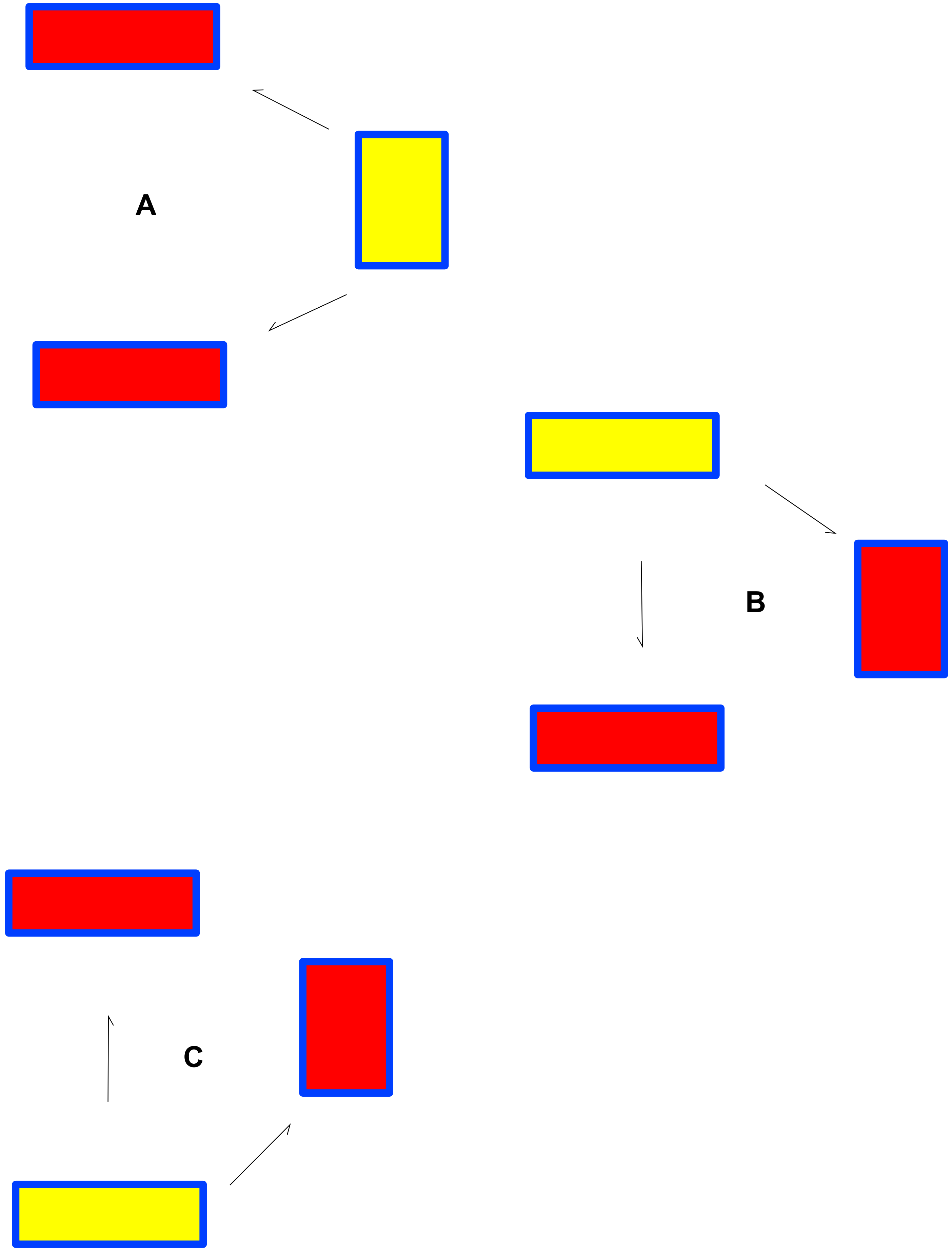

### Fig. S5

##### Agua Doce

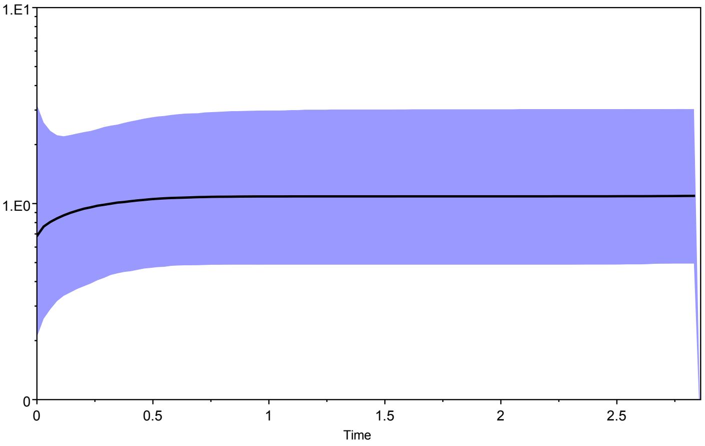


##### Boa Esperanca

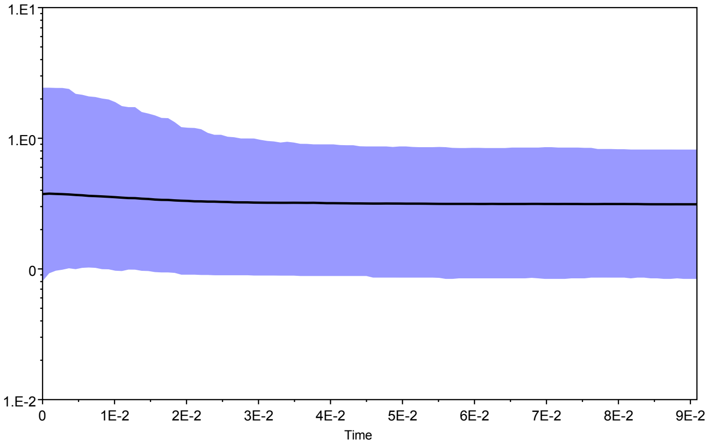


##### Canto

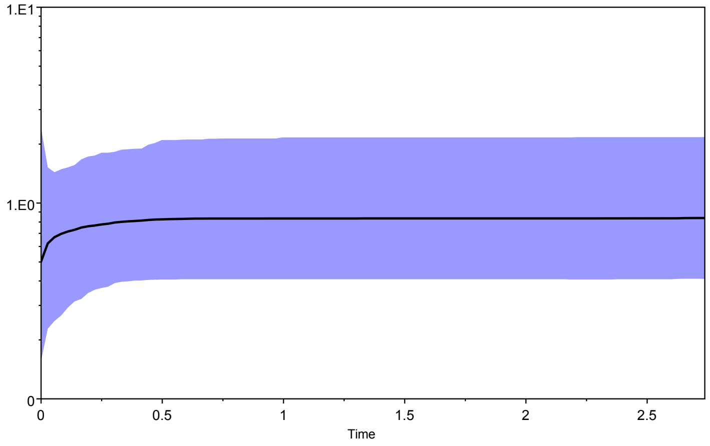


##### Curral Velho

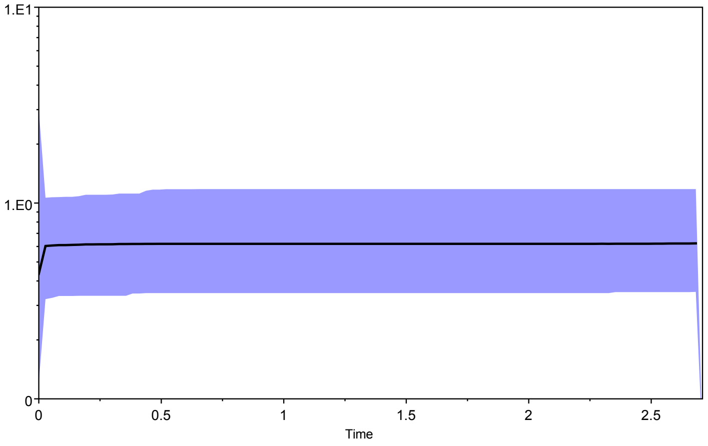


##### Norte

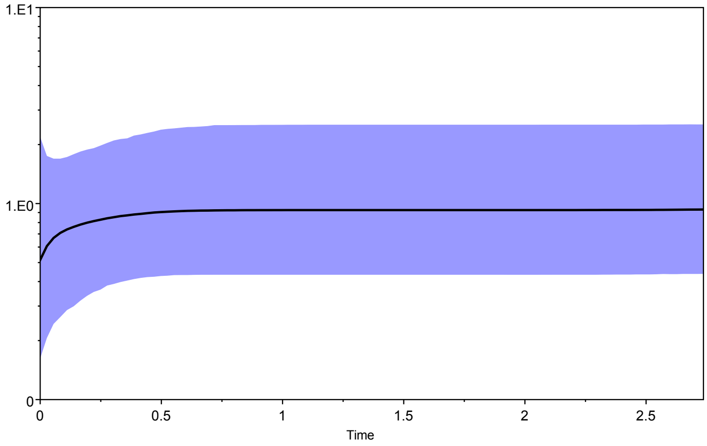


##### Ponta Pesqueira

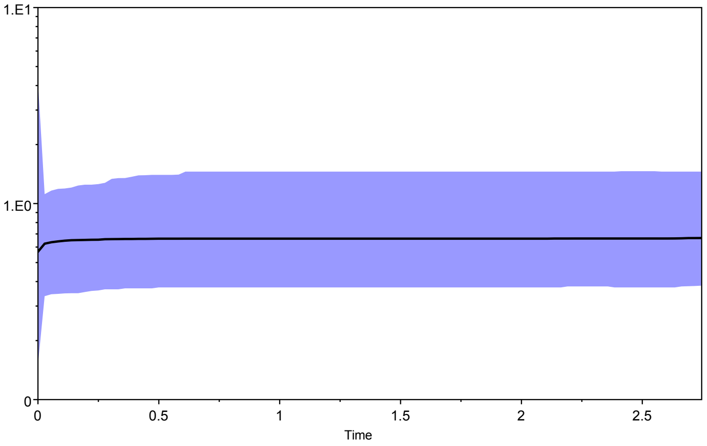


##### Lacacao

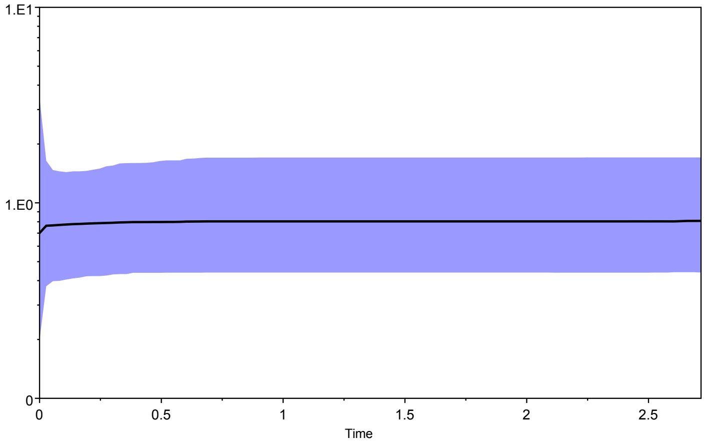
